## Appendix - Accession numbers for "Quantifying transmission dynamics of acute hepatitis C virus infections in a heterogeneous population using sequence data"

**Quantifying transmission dynamics of acute hepatitis C virus infections in men having sex with men using phylodynamics**

Gonché Danesh, Victor Virlogeux, Christophe Ramière, Caroline Charre, Laurent Cotte, Samuel Alizon

### Appendix

**Table S1.** GenBank accession numbers of sequences used to infer the viral phylogeny. Each Accession number is associated to a host type ('classical' or 'new') and a sampling date.

| Accession number | Host type | Sampling date |
|---|---|---|
| KY928360 | classical | 31/07/2014 |
| KY928361 | classical | 24/09/2014 |
| KY928362 | classical | 06/10/2014 |
| KY928363 | classical | 17/10/2014 |
| KY928364 | classical | 20/11/2014 |
| KY928365 | classical | 01/12/2014 |
| KY928366 | classical | 05/01/2015 |
| KY928367 | classical | 13/01/2015 |
| KY928368 | classical | 16/01/2015 |
| KY928369 | classical | 16/01/2015 |
| KY928370 | classical | 20/01/2015 |
| KY928371 | classical | 30/01/2015 |
| KY928372 | classical | 04/02/2015 |
| KY928373 | classical | 06/02/2015 |
| KY928374 | classical | 10/02/2015 |
| KY928375 | classical | 11/02/2015 |
| KY928376 | classical | 13/02/2015 |
| KY928377 | classical | 17/02/2015 |
| KY928378 | classical | 17/02/2015 |
| KY928379 | classical | 18/02/2015 |
| KY928380 | classical | 23/02/2015 |
| KY928381 | classical | 25/02/2015 |
| KY928382 | classical | 02/03/2015 |
| KY928383 | classical | 04/03/2015 |
| KY928384 | classical | 05/03/2015 |
| KY928385 | classical | 09/03/2015 |
| KY928386 | classical | 16/03/2015 |
| KY928387 | classical | 18/03/2015 |

| Accession number | Host type | Sampling date |
|---|---|---|
| KY928388 | classical | 27/03/2015 |
| KY928389 | classical | 30/03/2015 |
| MH885712 | classical | 31/03/2015 |
| MH885713 | classical | 31/03/2015 |
| MH885714 | classical | 10/04/2015 |
| MH885715 | classical | 14/04/2015 |
| MH885716 | classical | 15/04/2015 |
| MH885717 | classical | 21/04/2015 |
| MH885718 | classical | 28/04/2015 |
| MH885719 | classical | 30/04/2015 |
| MH885720 | classical | 30/04/2015 |
| MH885721 | classical | 30/04/2015 |
| MH885722 | classical | 07/05/2015 |
| MH885723 | classical | 20/05/2015 |
| MH885724 | classical | 26/05/2015 |
| MH885725 | classical | 28/05/2015 |
| MH885726 | classical | 03/06/2015 |
| MH885727 | classical | 05/06/2015 |
| MH885728 | classical | 08/06/2015 |
| MH885729 | classical | 09/06/2015 |
| MH885730 | classical | 10/06/2015 |
| MH885731 | classical | 11/06/2015 |
| MH885732 | classical | 12/06/2015 |
| MH885733 | classical | 12/06/2015 |
| MH885734 | classical | 17/06/2015 |
| MH885735 | classical | 18/06/2015 |
| MH885736 | classical | 19/06/2015 |
| MH885737 | classical | 03/07/2015 |
| MH885738 | classical | 08/07/2015 |
| MH885739 | classical | 22/07/2015 |
| MH885740 | classical | 24/07/2015 |
| MH885741 | classical | 28/07/2015 |
| MH885742 | classical | 29/07/2015 |
| MH885743 | classical | 30/07/2015 |
| MH885744 | classical | 12/08/2015 |
| MH885745 | classical | 17/08/2015 |
| MH885746 | classical | 17/08/2015 |
| MH885747 | classical | 18/08/2015 |
| MH885748 | classical | 21/08/2015 |
| MH885749 | classical | 24/08/2015 |

| Accession number | Host type | Sampling date |
|---|---|---|
| MH885750 | classical | 27/08/2015 |
| MH885751 | classical | 07/09/2015 |
| MH885752 | classical | 08/09/2015 |
| MH885753 | classical | 08/09/2015 |
| MH885754 | classical | 09/09/2015 |
| MH885755 | classical | 10/09/2015 |
| MH885756 | classical | 16/09/2015 |
| MH885757 | classical | 17/09/2015 |
| MH885758 | classical | 22/09/2015 |
| MH885759 | classical | 29/09/2015 |
| MH885760 | classical | 29/09/2015 |
| MH885761 | classical | 06/10/2015 |
| MH885762 | classical | 12/10/2015 |
| MH885763 | classical | 19/10/2015 |
| MH885764 | classical | 20/10/2015 |
| MH885765 | classical | 21/10/2015 |
| MT108308 | classical | 28/10/2015 |
| MT108309 | classical | 28/10/2015 |
| MT108310 | classical | 06/11/2015 |
| MT108311 | classical | 20/11/2015 |
| MT108312 | classical | 25/11/2015 |
| MT108313 | classical | 02/12/2015 |
| MT108314 | classical | 04/12/2015 |
| MT108315 | classical | 07/12/2015 |
| MT108316 | classical | 11/12/2015 |
| MT108317 | classical | 11/12/2015 |
| MT108318 | classical | 14/12/2015 |
| MT108319 | classical | 30/12/2015 |
| MT108320 | classical | 04/01/2016 |
| MT108321 | classical | 05/01/2016 |
| MT108322 | classical | 05/01/2016 |
| MT108323 | classical | 08/01/2016 |
| MT108324 | classical | 15/01/2016 |
| MT108325 | classical | 15/01/2016 |
| MT108326 | classical | 19/01/2016 |
| MT108327 | classical | 03/02/2016 |
| MT108328 | classical | 05/02/2016 |
| MT108329 | classical | 05/02/2016 |
| MT108330 | classical | 16/02/2016 |
| MT108331 | classical | 16/02/2016 |

| Accession number | Host type | Sampling date |
|---|---|---|
| MT108332 | classical | 19/02/2016 |
| MT108333 | classical | 26/02/2016 |
| MT108334 | classical | 07/03/2016 |
| MT108335 | classical | 07/03/2016 |
| MT108336 | classical | 30/03/2016 |
| MT108337 | classical | 01/04/2016 |
| MT108338 | classical | 01/04/2016 |
| MT108339 | classical | 07/04/2016 |
| MT108340 | classical | 18/04/2016 |
| MT108341 | classical | 25/04/2016 |
| MT108342 | classical | 26/04/2016 |
| MT108343 | classical | 19/05/2016 |
| MT108344 | classical | 25/05/2016 |
| MT108345 | classical | 26/05/2016 |
| MT108346 | classical | 27/05/2016 |
| MT108347 | classical | 03/06/2016 |
| MT108348 | classical | 06/06/2016 |
| MT108349 | classical | 07/06/2016 |
| MT108350 | classical | 13/06/2016 |
| MT108351 | classical | 21/06/2016 |
| MT108352 | classical | 01/07/2016 |
| MT108353 | classical | 04/07/2016 |
| MT108354 | classical | 07/09/2016 |
| MT108355 | classical | 08/09/2016 |
| MT108356 | classical | 06/01/2017 |
| MT108357 | classical | 08/02/2017 |
| MT108358 | classical | 01/06/2017 |
| MT108359 | classical | 05/09/2017 |
| MT108360 | classical | 22/09/2017 |
| MT108361 | classical | 10/10/2017 |
| MT108362 | classical | 24/04/2018 |
| MT108363 | classical | 24/04/2018 |
| MT108364 | classical | 02/05/2018 |
| MT108365 | classical | 02/05/2018 |
| MT108366 | classical | 02/05/2018 |
| MT108367 | classical | 03/05/2018 |
| MT108368 | classical | 24/05/2018 |
| MH885654 | new | 27/07/2011 |
| MH885655 | new | 21/01/2013 |
| KY928329 | new | 15/02/2013 |

| **Accession number** | **Host type** | **Sampling date** |
|---|---|---|
| KY928344 | new | 23/05/2013 |
| KY928348 | new | 24/05/2013 |
| KY928322 | new | 16/12/2013 |
| MH885656 | new | 18/02/2014 |
| MH885657 | new | 04/04/2014 |
| KY928330 | new | 22/09/2014 |
| MH885658 | new | 22/10/2014 |
| KY928355 | new | 13/02/2015 |
| MH885659 | new | 26/05/2015 |
| KY928356 | new | 27/05/2015 |
| MH885660 | new | 10/06/2015 |
| KY928352 | new | 12/06/2015 |
| MH885661 | new | 12/06/2015 |
| MH885662 | new | 15/06/2015 |
| KY928313 | new | 24/07/2015 |
| KY928314 | new | 06/08/2015 |
| MH885663 | new | 16/09/2015 |
| MH885664 | new | 05/10/2015 |
| KY928336 | new | 14/10/2015 |
| KY928338 | new | 05/11/2015 |
| KY928350 | new | 16/11/2015 |
| MH885665 | new | 29/12/2015 |
| MH885666 | new | 30/12/2015 |
| MH885667 | new | 05/01/2016 |
| MH885668 | new | 03/02/2016 |
| MH885669 | new | 10/02/2016 |
| MH885670 | new | 03/03/2016 |
| KY928357 | new | 23/03/2016 |
| KY928341 | new | 24/03/2016 |
| MH885671 | new | 06/04/2016 |
| KY928354 | new | 07/04/2016 |
| KY928311 | new | 15/04/2016 |
| KY928335 | new | 31/05/2016 |
| KY928346 | new | 02/06/2016 |
| MH885672 | new | 13/06/2016 |
| MH885673 | new | 05/07/2016 |
| KY928349 | new | 03/08/2016 |
| KY928331 | new | 01/09/2016 |
| MH885674 | new | 14/09/2016 |
| KY928324 | new | 21/09/2016 |

| Accession number | Host type | Sampling date |
|---|---|---|
| KY928353 | new | 19/10/2016 |
| MH885675 | new | 04/11/2016 |
| KY928351 | new | 30/11/2016 |
| KY928343 | new | 23/01/2017 |
| MH885676 | new | 10/02/2017 |
| KY928325 | new | 22/03/2017 |
| MH885677 | new | 27/03/2017 |
| KY928333 | new | 05/04/2017 |
| KY928345 | new | 03/05/2017 |
| KY928315 | new | 27/06/2017 |
| MH885693 | new | 10/07/2017 |
| MH885695 | new | 19/07/2017 |
| MH885697 | new | 24/07/2017 |
| MH885698 | new | 25/07/2017 |
| MH885699 | new | 16/08/2017 |
| MH885700 | new | 17/08/2017 |
| MH885701 | new | 21/08/2017 |
| MH885702 | new | 30/08/2017 |
| MH885703 | new | 27/07/2017 |
| MH885704 | new | 03/11/2017 |
| MH885705 | new | 14/11/2017 |
| MH885707 | new | 27/11/2017 |
| MH885711 | new | 20/12/2017 |
| MT108306 | new | 22/11/2017 |
| MT108307 | new | 27/11/2017 |
